# Namdinator - Automatic Molecular Dynamics flexible fitting of structural models into cryo-EM and crystallography experimental maps

**DOI:** 10.1101/501197

**Authors:** Rune Thomas Kidmose, Jonathan Juhl, Poul Nissen, Thomas Boesen, Jesper Lykkegaard Karlsen, Bjørn Panyella Pedersen

## Abstract

Model building into experimental maps is a key element of structural biology, but can be both time consuming and error-prone. Here we present Namdinator, an easy-to-use tool that enables the user to run a Molecular Dynamics Flexible Fitting (MDFF) simulation in an automated manner through a pipeline system. Namdinator will modify an atomic model to fit within cryo-EM or crystallography density maps, and can be used advantageously for both the initial fitting of models, and for a geometrical optimization step to correct outliers, clashes and other model problems. We have benchmarked Namdinator against 39 deposited models and maps from cryo-EM and observe model improvements in 34 of these cases (87%). Clashes between atoms were reduced, and model-to-map fit and overall model geometry were improved, in several cases substantially. We show that Namdinator is able to model large scale conformational changes compared to the starting model. Namdinator is a fast and easy way to create suitable initial models for both cryo-EM and crystallography. It can fix model errors in the final steps of model building, and is usable for structural model builders at all skill levels. Namdinator is available as a web service (https://namdinator.au.dk), or can be run locally as a command-line tool.

**Synopsis:** A pipeline tool called Namdinator is presented that enables the user to run a Molecular Dynamics Flexible Fitting (MDFF) simulation in a fully automated manner, both online and locally. This provides a fast and easy way to create suitable initial models for both cryo-EM and crystallography and help fix errors in the final steps of model building.

## 1. INTRODUCTION

In recent years, major technical advances in the cryo-EM field have resulted in an increasing number of cryo-EM density maps being deposited (Kühlbrandt, 2014). With the growing number of maps follows an increasing demand for better and easier methods for fitting atomic models. An analysis of models fitted to cryo-EM maps has previously indicated the presence of notable problems in almost all of the selected structures (Wlodawer *et al*., 2017). In 2008, the powerful Molecular Dynamics Flexible Fitting (MDFF) method was presented. MDFF can fit a model in a flexible manner into a cryo-EM density map, using molecular dynamic simulations (Trabuco *et al*., 2008). Implementing MDFF as a standard tool for model building involves a steep learning curve which, among other things, consist of preparing and restraining the input model for the simulation, converting the map to a potential field and finally setting up and running the simulation. Namdinator was developed as an user-friendly automated MDFF pipeline to accommodate the increase in available cryo-EM reconstructions of macromolecular structures.

Through testing and benchmarking we demonstrate that Namdinator is able to obtain excellent fits of models to cryo-EM maps with no manual intervention needed. Namdinator does not only optimize the fit to the map, but it also improves the quality and geometry of the fitted model. Besides cryo-EM maps, Namdinator is also useful for crystallographic electron density maps. As such Namdinator speeds up model building and improves the quality of the final model for both crystallographic and cryo-EM data.

## 2. EXPERIMENTAL

### 2.1. Namdinator framework

A flowchart of Namdinator is presented in Figure 1. To run Namdinator, a model in standard Protein Data Bank (PDB) file format and a map is needed. The map can be a cryo-EM density map or a crystallographic map in either mrc or ccp4 format, and it should be in a P1 space group (default for EM). The initial model can be a homology model or a model of the target protein in a different state/conformation if desired. The model should be roughly docked into the map before using Namdinator. Initial docking can be done manually or by rigid body fitting programs such as COLORES (Chacón & Wriggers, 2002) or Chimera (Pettersen *et al*., 2004).

**Figure 1.**
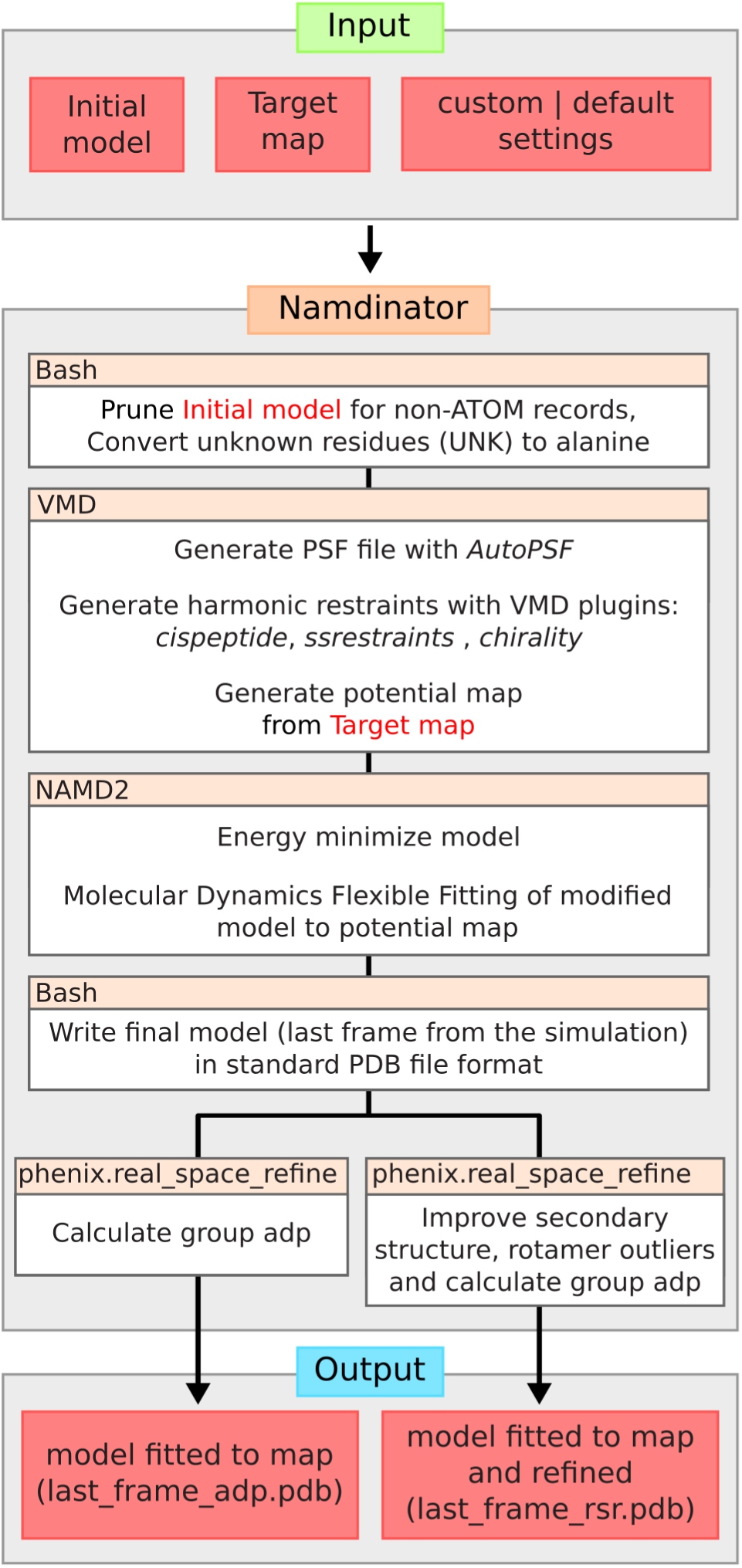
Flowchart for Namdinator. As input the user must provide an initial model and a target map with its corresponding resolution. A number of settings can be set or default values can be used. Namdinator then modifies the initial model and runs a NAMD2 simulation using MDFF. After the run, phenix.real_space_refine will be run on the output. Either calculating only group ADP or doing real space refinement and group ADP calculation. The latter, in particular, will help improve Ramachandran plot outliers (backbone/secondary structure improvements), and Rotamer outliers.

Within Namdinator the input PDB file is prepared for the MDFF simulation using the VMD-plugins MDFF and AutoPSF (Humphrey *et al*., 1996). AutoPSF uses an internal set of standard settings and topology files to generate dynamics-ready PDB and Protein Structure File (PSF) files. While AutoPSF allows for a streamlined process, it is prone to failure when encountering non-standard atoms in a PDB file. Therefore, all non-ATOM records are removed, meaning that metal ions, water molecules, ligands etc. will not be included in the simulation nor in the output PDB files, and must be manually reinserted if so required. All residues designated UNK within the input PDB file are converted to alanine residues. If the starting model is pruned for side chains but maintains sequence information, a corresponding full atomic model is generated and used during the simulation, which means the output will also be full atomic models.

During the MDFF simulation in NAMD2, harmonic restraints are applied to the input model in order to prevent over-fitting and structural distortion, and to preserve the stereo chemical quality of the model. The simulation is run using the CHARMM36 force field for all-atom systems in vacuum. *In vacuo* MDFF simulation leads to a substantial speed up (up to 7 fold) in calculation time compared to Generalized Born Implicit Solvent (GBIS) MDFF simulations (Tanner *et al*., 2011). Namdinator also supports GBIS simulations, which do yield marginally better results judged from the quality of the output models and obtained fit, but at the cost of a substantial increase in running time. After the MDFF simulation, the last frame of the resulting trajectory is exported as a PDB file, all hydrogen atoms are pruned and standard PDB atom and residue naming format is enforced. This PDB is then processed in two parallel setups through phenix.real_space_refine. In the first, only group Atomic Displacement Factors (ADP) are calculated, and the user can then directly use this output model. In the second setup, the model will be both have ADP calculated and be refined in real space (with settings: Secondary Structure, Rotamer, Ramachandran and C-beta deviation restraints, global minimization, Group ADP-refinement, 5 macro cycles) (Afonine *et al*., 2013). We find that this second setup is highly beneficial for cryo-EM models in order to reduce the number of Rotamer and Ramachandran plot outliers as well as C-beta deviations, as these parameters are not restrained during simulation. Crystallographic models, being subject to further crystallographic refinement downstream, does not require this step.

### 2.2. Output validation in Namdinator

Namdinator performs a validation check of both the input PDB file (After group ADP calculation) and the output PDB file from the simulation and from the phenix.real_space_refine run. For validation Namdinator uses the programs phenix.ramalyse, phenix.clashscore, phenix.rotalyse and phenix.cbetadev (Adams *et al*., 2010). A summary of the validation metrics for each PDB file is displayed in a table at the end of the simulation. Additionally, the RosettaCommons package is used to score the PDB files both as whole models and for individual residues (Alford *et al*., 2017). The Rosetta score for the models are included with the phenix validation metrics, whereas the top-10 flagged residues in each model are reported in a separate table at the end of the run and can readily be assessed for e.g. atom clashes. Together the two tables provide the user with a convenient overview to analyze and compare the models. To evaluate how the models fit to the input map, cross-correlation coefficients (CC) are calculated using phenix.map_model_cc, using the stated map resolution of the input map.

The web-service will load the input as well as the output models in an online 3D viewer (NGL viewer)(Rose *et al*., 2016) together with the target map, for rapid and user-friendly manual inspection of the result.

### 2.3. Namdinator system setup

Namdinator’s current third party software requirements are VMD v.1.93 with the plugins: MDFF v.0.5, ssrestraints v.1.1, cispeptides v.1.3, chirality v.1.3, Autopsf v.1.6 and multiplot v.1.7. All simulations are run within the CUDA optimized version of NAMD2 (2.12) (Phillips *et al*., 2005) using the CHARMM36 Force field (Huang & MacKerell, 2013) for all-atom systems in vacuum. MDFF applies default parameters (step size: 1 fs, force scaling (G scale): 0.3 kcal/mol [adjustable], temperature 300K [adjustable] using Langevin thermostat coupled to all non-hydrogen atoms with a damping coefficient of 5 ps-1, bonded interactions calculated every 1 fs, nonbonded interactions (cutoff 10 Å) calculated every 2 fs (Trabuco *et al*., 2009)). G-scale is a measure of how hard the target map pulls on the starting model. Temperature controls how easy it will be for atoms to move. For hard cases it can be beneficial to adjust one or both of these parameters from their default, as well as increasing the number of simulation steps.

Namdinator further depends on the phenix package for group ADP and real space refinement and validation together with the rosettaCommons software package. phenix.real_space_refine requires the presence of a CRYST1 record in the header of the input PDB file. If no CRYST1 record is present, Namdinator will append a standard CRYST1 record with unit cell dimensions of 1.00 for all three axes, 90 degrees for all three angles and P1 space group. During the phenix.real_space_refine run, the unit cell dimensions will be updated with that of the box size of the input map. Namdinator uses GPU accelerated calculation within NAMD2, and a CUDA-capable video card is needed for local installations. Additionally, as NAMD2 does not offload all of the calculations to the GPU, a high powered multi core CPU is recommended. A description of the specific flags that control Namdinator are listed in the command-line version of the program. The web-service will allow the user to run Namdinator and adjust the above settings within reasonable predefined ranges.

## 3. RESULTS AND DISCUSSION

### 3.1. Benchmarking using cryo-EM models and maps

To benchmark Namdinator, 39 randomly selected map and model pairs deposited to the Electron Microscopy Data Bank (EMDB) and PDB was run through Namdinator using the resolution stated on their EMDB entry pages. Five of the PDB files were manually modified to allow Namdinator to run: Two to contain unique single-character, case-insensitive chain-IDs (5GW5 and 5H64). One to have only positive residue numbering (6B44). One was a multi-model PDB, that was reduced to a single model (5ND7), and one was expanded to have its biological molecule generated (3J9C) from the deposited model. No atom coordinates were altered before use as input for Namdinator and no maps were filtered or modified in any manner. All test cases were run using default Namdinator settings. To simplify the comparison of input to output we have focused on 3 global key statistics. Cross Correlation (to gauge the fit between model and map), number of clashes (to gauge the chemical environment of the individual atoms of the models), and Ramachandran plot outliers (to gauge overall protein geometry).

Overall, Namdinator improved the test models (Fig. 2, Supplementary Fig. 1 and Supplementary Table 1). The CC improved in 22 out of the 39 test cases. It was unchanged in 11 cases (defined as +/- 1%), and deteriorated in 6 cases. The Clash score was improved in 17 cases, 18 cased had a similar clash score (defined as +/- 5), and in 4 cases the clash score deteriorated. The number of Ramachandran outliers was reduced in 23 of the cases, identical in 12 of the cases, and increased in 4 test cases.

**Figure 2.**
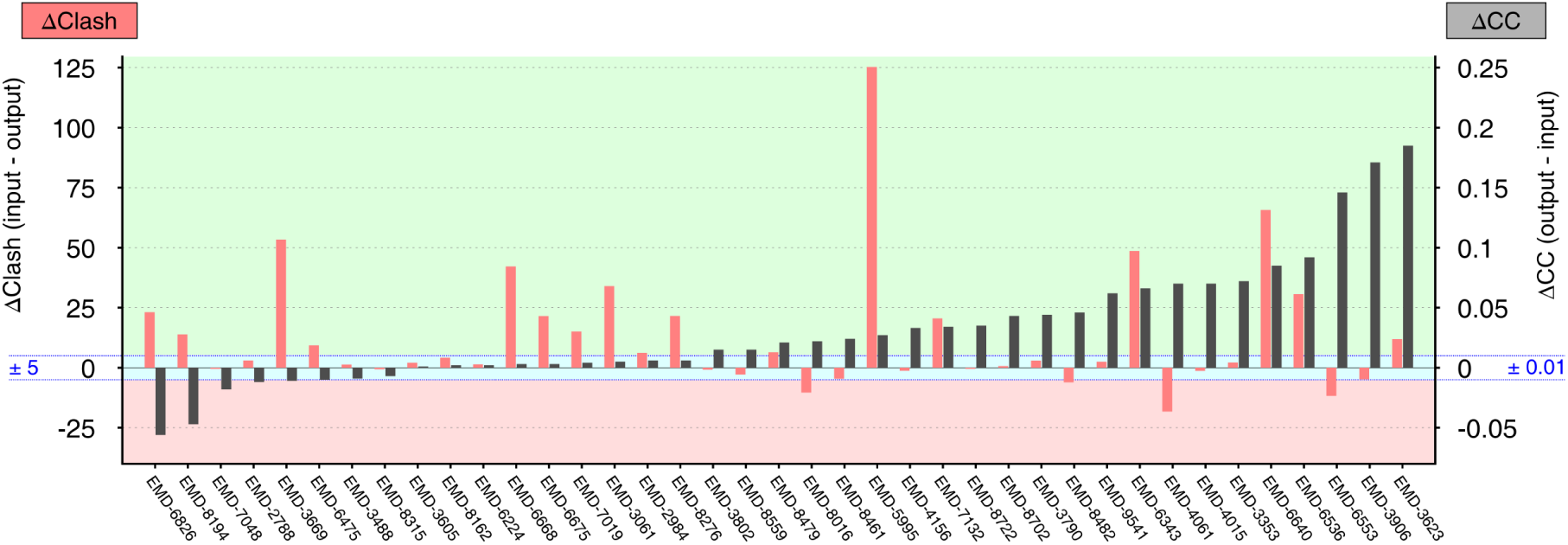
Benchmark of Namdinator with 39 deposited Cryo-EM structures and maps. Plot of global quality parameters; delta clash score (pink bars) and delta CC (grey bars). The individual EMDB entry ID’s are listed along the X-axis. The delta clash score is calculated by subtracting the clash score of the final model from the clash score of the input model. The delta CC value is calculated by subtracting the cross-correlation coefficient between input model and map from the cross-correlation coefficient between final model and map. In both cases a positive value indicates that the output from Namdinator was improved compared to the input. The green shade indicates model improvement through Namdinator, the red shade indicate model deterioration, and the blue shade a comparable quality model (+/5 clash or +/- 0.01 CC).

In total, of the 39 test cases, 34 models were improved, some substantially on at least one of the three key statistics. In the majority of test cases where statistical scores deteriorated after Namdinator, the starting models were not full-atom models. As Namdinator convert these to full atom models, the change in most cases can be attributed to this conversion. The five cases where all three key statistics did not improve (6B44, 5NI1, 5SY1, 5N9Y, 3J9C) were relatively high resolution (3.9 - 2.9 Å) full-atom models, where we would expect Namdinator to have the least impact.

The use of general statistics to evaluate quality can hide pronounced local improvements. Visual inspection showed further quality improvements that are not easily quantified. One example was the Peptide Loading Complex (PLC), using the map EMDB-3906 and its associated model 6ENY, which was deposited as a poly-alanine model (Blees *et al*., 2017). The initial clash score of 6ENY is 12.9, with 33 Ramachandran outliers. The CC between the deposited model and the map is 56.2%. The output model from Namdinator, which now is a full atomic model, obtains a clash score of 17.7 with 2 Ramachandran outliers, while the CC increases to 73.6%. Since 6ENY is a poly-alanine model the input model clashes originate from CB and/or backbone atoms only and indicates severe backbone clashes in the model. By manually inspecting the residues with the highest individual rosetta scores, as listed in the validation table output from Namdinator, several backbone clashes were readily identified in the input model, which had been fixed by Namdinator (Fig. 3).

**Figure 3.**
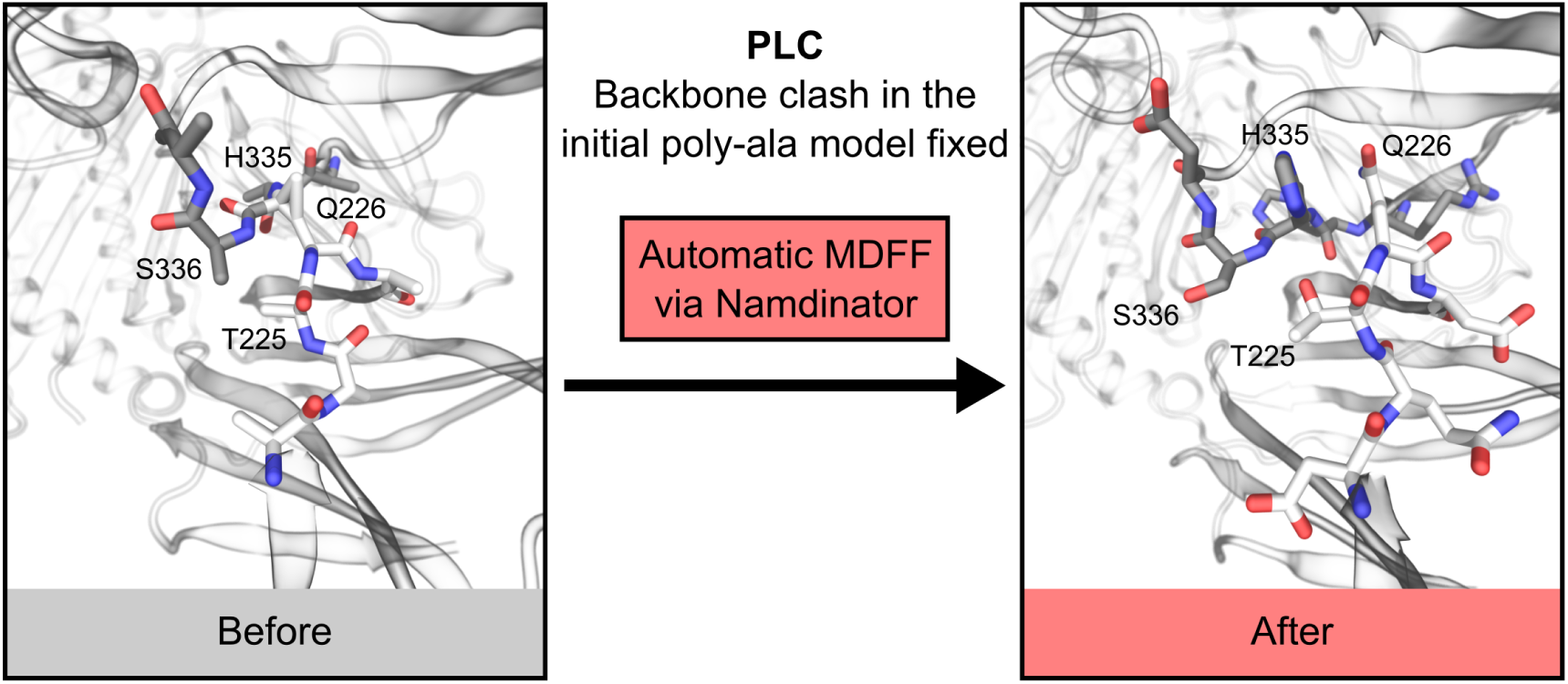
Example of local improvements through Namdinator. A severe backbone clash was present in the deposited poly-Ala model of the peptide loading complex PLC (6ENY), between residue Gln226 in chain F and His335 in chain C. Namdinator fixed this clash, and by looking at the Namdinator log file is was simple to identify the problematic regions in the input model.

Judged on global statistics, Namdinator improved 87% of the test cases. Several models revealed quite extensive improvements, where severe backbone clashes and unnatural and strained conformations were reduced. All of these are model problems that also can be addressed manually, but clearly an automated method like Namdinator is of great assistance to create higher quality models based on objective criteria and global targets.

During testing of deposited maps and models, one case was an obvious outlier with a huge CC increase (EMD-3765/5O9G) (Farnung *et al*., 2017). Upon inspection, it was observed that the model was systematically shifted in one direction relative to the map. The systematic shift had occurred during the data deposition, without the authors’ knowledge (Farnung, personal communication). Namdinator caught this shift and moved the model back into density within the first 1000 simulation steps. This case is omitted from the final list of examples, but it illustrates the use of Namdinator as a rigid body fitting tool even for models that are systematically shifted relative to their density.

### 3.2. Large-scale movements and conformational changes

We tested Namdinator’s ability to fit models, when large parts of the starting model were placed outside the target density. MDFF has been used successfully for fitting one conformation of a protein into the map of a different conformation of the same protein, and Namdinator should be able to handle these scenarios with no or minimal intervention.

As a classic MDFF example the model of the *E.coli* adenylate kinase (Lou & Cukier, 2006), locked in a catalytic transition state by an inhibitor (1AKE)(Müller & Schulz, 1992) was fitted into a 5 Å simulated density of an apo-state of the adenylate kinase (4AKE)(Müller *et al*., 1996) using Namdinator. After 60,000 simulation steps a very good fit was obtained and the CC had increased from 39.0% to 71.3%. (Fig. 4A and Supp. Movie 1). This shows that Namdinator can model large movements between conformational states with a noise-free simulated map (Derived from a PDB model) as the test case.

**Figure 4.**
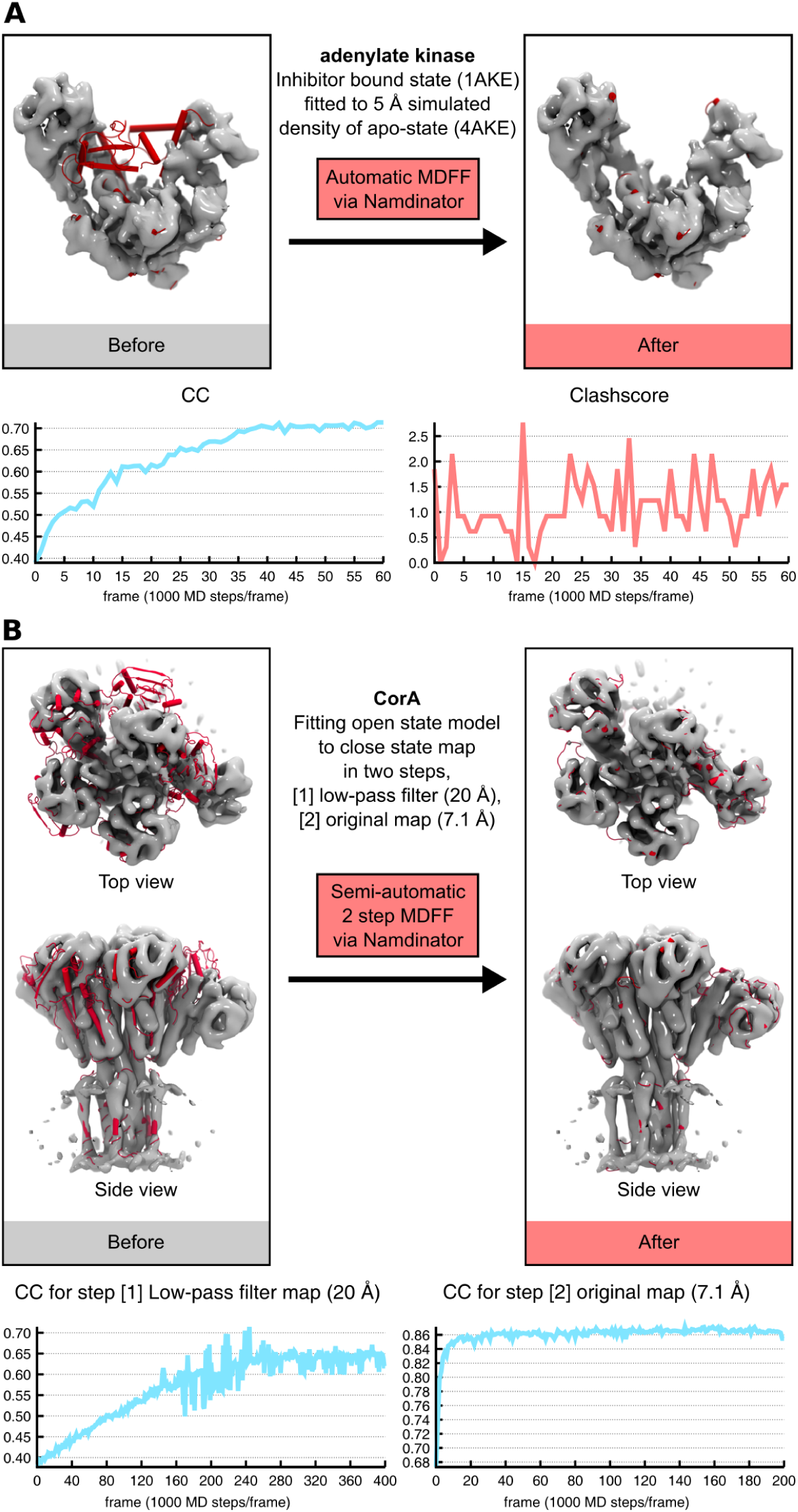
Large-scale movements of models into density. (A). Fitting of *E.coli* Adenylate Kinase (1AKE) into a 5 Å simulated density of the adenylate kinase in a different conformation (4AKE) using Namdinator with plot of the CC values and clash score for every 1,000 step from Namdinator. Default settings were used, except the simulation steps which were increased from 20.000 to 60.000 to reach convergence. (**B**) Fitting the open conformation of the magnesium channel CorA into the density (EMD-6553) of its closed state using Namdinator. The initial model (red cartoon) rigid-fitted to EMD-6553 (grey surface), that was used as input for a two-step Namdinator procedure. A plot of the CC values for every 1,000 step from both sequential Namdinator runs are shown.

Namdinator can also handle more challenging cases, such as noisy maps. We used the magnesium channel CorA as a test case (Matthies *et al*., 2016), where three different conformations of CorA have been determined by cryo-EM; a closed conformation at 3.8 Å and two open conformations both at 7.1 Å. During transition from closed to open conformations, CorA changes from being a 5-fold symmetric structure to an asymmetric structure where 4 of 5 subunits are being displaced between 10 to 25 Å and intra subunits undergoes hinge-bending motions with up to 35 degrees (Matthies *et al*., 2016). To perform the fit, the closed conformation model (3JCF/EMD-6551) was first rigid-body fitted into the original 7.1 Å cryo-EM density map of the open state conformation (3JCH/EMD-6553) using COLORES. An initial run revealed that a default Namdinator setup lead to domains and secondary structure elements becoming stuck at intermediary positions during the fit. To solve this, the map was low passed filtered to 20 Å, before being piped into Namdinator. Default Namdinator settings was used except for the G-scale which was decreased from the default of 0.3 kcal/mol to 0.05 kcal/mol while the number of steps was increased to 400,000 steps. The output of an initial run with the low pass filtered map was then used as input for a second Namdinator run fitting into the original unfiltered 7.1 Å map where again default settings was used except for the G-scale which was increased to 5 kcal/mol and the number of steps which was increased to 200,000. With this two-step procedure, Namdinator successfully fitted the 3JCF model into the density of the open state conformation of CorA. The final Namdinator result was compared with the deposited model for the open state of CorA. The RMSD of the two models was 1.6 Å, i.e. an excellent fit within the experimental error of the models at 7.1 Å resolution, but with little manual intervention (Fig. 4B, Supp. Movie 2).

### 3.3. Crystallographic models and maps

Namdinator is designed to handle models and maps from crystallography as well as cryo-EM. We tested 10 low resolution crystal structures along with their corresponding map coefficients downloaded from the PDB. Namdinator was not able to optimize these structures in a consistent way, i.e. unlike what was observed for the cryo-EM models analyzed. This is perhaps expected as the crystallographic models are heavily restrained by the crystallographic refinement and the deposited models have presumably been optimized already to maximize the fit between observations and model.

However, Namdinator is still very useful for initial model building in crystallographic electron density maps of poor quality and can save the user considerable time during the initial steps of model building. We find a clear benefit of using Namdinator in crystallography in the early steps of model building based on poorly fitting starting models. In-house Namdinator has become our go-to tool for this purpose.

## 4. CONCLUDING REMARKS

Namdinator is an automatic and user-friendly pipeline for flexible fitting and geometrical optimization of a given model to a given map. Running time is fast and we find that as a versatile tool it can efficiently facilitate model building both in cryo-EM and crystallography. Importantly, Namdinator enables higher quality depositions compared to the current state-of-the-art (Wlodawer *et al*., 2017). Clearly, special cases will occur, but as demonstrated here, a default Namdinator run will improve model geometry, while maintaining or improving the fit to the map in almost all cases, and for difficult cases simple adjustments of parameters and procedures will support the application.

Namdinator is a fast way to generate initial plausible models and to correct geometrical errors in the final steps and is a valuable tool in the challenging process of constructing structural models. In the end there is no substitute for human interaction during model building, and Namdinator (and similar tools) should not be seen as a replacement for a final and careful manual inspection of a model, but solely as a tool to assist and speed up this process.

*Namdinator is released under the GNU General Public License version 3 (GNU GPLv3) and is freely available as a linux command-line tool on github (https://github.com/namdinator) or by contact to the corresponding author (JLK). A web-service is available at* https://namdinator.au.dk.

## Acknowledgements

We are grateful to the Centre for Structural Biology for discussions and beta-testing. This work was supported in part by funding from the European Research Council (grant agreement No. 637372), the Danish Council for Independent Research (grant agreement no. DFF-4002-00052) as well as an AIAS fellowship to B.P.P.

## Supplementary Material

**Supplementary Figure 1.**
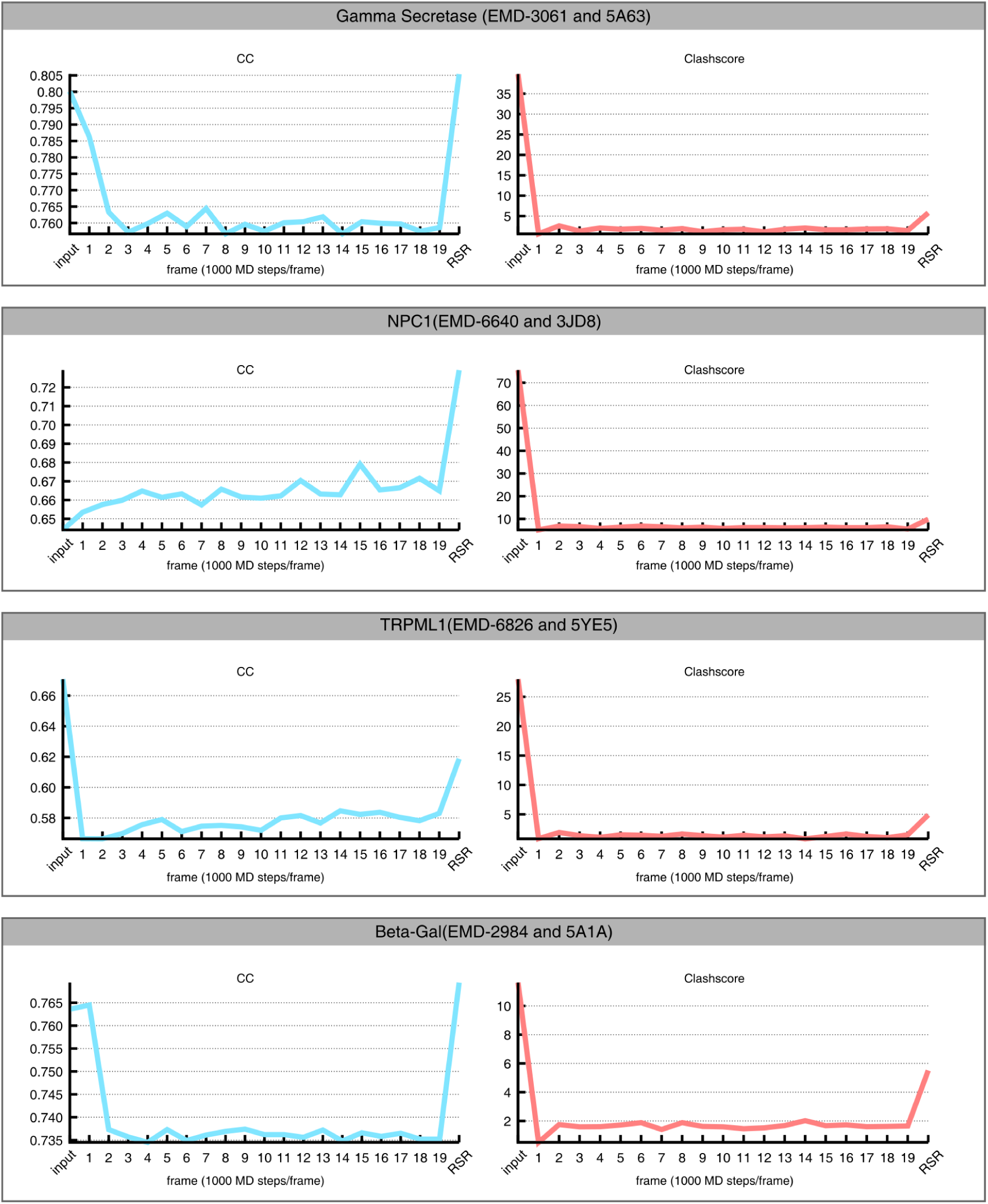
Selected benchmark test cases in detail. At every 1000th simulation step Namdinator exported the coordinates as a PDB file. The B-factors of the output frames were refined in phenix.real_space_refine using group ADP refinement only. Their clash score and cross correlation is plotted as a function of frame-number. The input model (input) was also subjected to group ADP refinement for CC calculation. The final output model from Namdinator after a default real space refinement run in phenix.real_space_refine (RSR) is also plotted. All runs used Namdinator default settings.

**Supplementary Table 1. Table of 39 test cases run through Namdinator.** Legend: FAT = Full atomic model; PA = Poly-Ala model; CA = backbone model; FAT/PA = Full atomic model mixed with Poly-Ala segments; LF = Last_frame_adp.pdb; LF_RSR = Last_frame_rsr.pdb. As discussed in the main text, when comparing the results, especially one caveat must be taken into account: Due to resolution-limits and other considerations, many of the tested input models were not deposited as full atomic models, but deposited as either backbone only, pruned to poly-Ala only or a mix of full atomic residues and either of the two former options. After Namdinator all models are now full atom models. The clash score is calculated by phenix.clashscore as follows: Hydrogens are removed, then all atomic overlaps worse than 0.4 Å are listed, and a global clash score is calculated for the entire model calculated as 1000 * (number of bad overlaps) / (number of atoms).

**Supplementary Movie 1.** Fitting of *E.coli* Adenylate Kinase (1AKE) into a 5 Å simulated density of the adenylate kinase in a different conformation (4AKE) using Namdinator.

**Supplementary Movie 2.** Two-step fitting of the closed conformation of CorA (3JCF) into the open conformation (EMD-6553).

