## Supplementary Table 1 for "Namdinator - Automatic Molecular Dynamics flexible fitting of structural models into cryo-EM and crystallography experimental maps"

| pdb | map | res. (Å) | model type | | Clashscore | Ramachandran plot Favored | Allowed | Outliers | C-beta deviations | Rotamer outliers | phenix CC | Cis-peptides | Rosetta score |
|---|---|---|---|---|---|---|---|---|---|---|---|---|---|
| 5ye5 | 6826 | 5.8 | CA/PA | **Input** | 28.01 | 1720 | 56 | 32 | 0 | 22.22% | 0.6750 | 24 | 80206 |
| | | | | **LF** | 1.85 | 1651 | 85 | 72 | 135 | 6.39% | 0.5887 | 23 | -7124 |
| | | | | **LF_RSR** | 4.90 | 1640 | 152 | 16 | 0 | 0.24% | 0.6187 | 20 | -8510 |
| 5k12 | 8194 | 1.8 | FAT | **Input** | 15.66 | 1674 | 66 | 0 | 0 | 0.82% | 0.8670 | 6 | -122915 |
| | | | | **LF** | 1.02 | 1652 | 74 | 14 | 68 | 2.53% | 0.8493 | 6 | -120715 |
| | | | | **LF_RSR** | 1.86 | 1716 | 24 | 0 | 0 | 0.41% | 0.8202 | 6 | -128126 |
| 6b44 | 7048 | 2.9 | FAT/PA | **Input** | 5.34 | 2528 | 326 | 0 | 0 | 0.43% | 0.7664 | 0 | -51090 |
| | | | | **LF** | 2.48 | 2496 | 266 | 92 | 114 | 4.30% | 0.6833 | 0 | -92158 |
| | | | | **LF_RSR** | 5.78 | 2617 | 236 | 1 | 1 | 0.13% | 0.7488 | 0 | -107532 |
| 4v1w | 2788 | 4.7 | FAT | **Input** | 7.78 | 3889 | 119 | 24 | 172 | 7.67% | 0.8161 | 0 | -18504 |
| | | | | **LF** | 1.17 | 3820 | 169 | 43 | 351 | 2.31% | 0.7736 | 0 | -21703 |
| | | | | **LF_RSR** | 4.77 | 3912 | 120 | 0 | 0 | 0.00% | 0.8040 | 0 | -24099 |
| 5np0 | 3669 | 5.7 | PA | **Input** | 67.09 | 4463 | 393 | 14 | 0 | 0.00% | 0.8063 | 0 | 748887 |
| | | | | **LF** | 10.23 | 4254 | 380 | 238 | 393 | 7.83% | 0.7697 | 0 | -40843 |
| | | | | **LF_RSR** | 13.74 | 4321 | 545 | 14 | 6 | 0.57% | 0.7949 | 0 | -54879 |
| 3jbr | 6475 | 4.2 | FAT/PA | **Input** | 18.27 | 2067 | 207 | 172 | 0 | 0.37% | 0.6931 | 25 | 63922 |
| | | | | **LF** | 3.34 | 2115 | 200 | 131 | 154 | 4.33% | 0.6106 | 25 | -10028 |
| | | | | **LF_RSR** | 9.01 | 2121 | 307 | 18 | 1 | 0.47% | 0.6831 | 25 | -11471 |
| 5ni1 | 3488 | 3.2 | FAT | **Input** | 2.97 | 539 | 27 | 0 | 0 | 0.43% | 0.8352 | 0 | -22631 |
| | | | | **LF** | 0.46 | 540 | 21 | 5 | 24 | 2.38% | 0.7858 | 0 | -20323 |
| | | | | **LF_RSR** | 1.71 | 548 | 18 | 0 | 0 | 0.00% | 0.8265 | 0 | -22862 |
| 5sy1 | 8315 | 3.9 | FAT | **Input** | 5.24 | 1341 | 103 | 2 | 0 | 0.08% | 0.7808 | 0 | -11632 |
| | | | | **LF** | 3.37 | 1336 | 90 | 20 | 62 | 2.66% | 0.7364 | 0 | -13686 |
| | | | | **LF_RSR** | 5.84 | 1372 | 72 | 2 | 0 | 0.00% | 0.7735 | 0 | -17823 |
| 5n9y | 3605 | 4.2 | FAT | **Input** | 12.67 | 1490 | 135 | 0 | 0 | 2.02% | 0.8259 | 0 | -12745 |
| | | | | **LF** | 1.72 | 1483 | 104 | 38 | 90 | 8.66% | 0.7904 | 0 | -11940 |
| | | | | **LF_RSR** | 10.68 | 1544 | 81 | 0 | 0 | 0.72% | 0.8266 | 0 | -13580 |
| 5jlf | 8162 | 3.6 | FAT/PA | **Input** | 8.79 | 2012 | 54 | 10 | 17 | 1.29% | 0.7302 | 5 | -30656 |
| | | | | **LF** | 1.42 | 1962 | 79 | 35 | 125 | 4.98% | 0.7044 | 5 | -28221 |
| | | | | **LF_RSR** | 4.63 | 1997 | 74 | 5 | 0 | 0.00% | 0.7317 | 5 | -29989 |
| 3j9c | 6224 | 2.9 | FAT | **Input** | 6.07 | 401 | 20 | 0 | 0 | 6.13% | 0.8286 | 1 | -18521 |
| | | | | **LF** | 0.76 | 390 | 27 | 4 | 7 | 3.47% | 0.7890 | 1 | -17624 |
| | | | | **LF_RSR** | 4.70 | 405 | 16 | 0 | 0 | 0.27% | 0.8304 | 1 | -19775 |
| 5h64 | 6668 | 4.4 | FAT/PA | **Input** | 51.29 | 5088 | 1098 | 154 | 0 | 0.80% | 0.7278 | 134 | 203740 |
| | | | | **LF** | 3.20 | 5057 | 961 | 322 | 455 | 4.41% | 0.7057 | 134 | -32771 |
| | | | | **LF_RSR** | 9.10 | 5606 | 685 | 49 | 1 | 0.39% | 0.7305 | 129 | -40825 |
| 5wq7 | 6675 | 3.0 | FAT | **Input** | 26.87 | 6630 | 285 | 45 | 15 | 0.49% | 0.8152 | 15 | -230152 |
| | | | | **LF** | 1.60 | 6492 | 373 | 95 | 317 | 4.12% | 0.7790 | 15 | -214396 |
| | | | | **LF_RSR** | 5.39 | 6701 | 259 | 0 | 0 | 0.00% | 0.8182 | 15 | -239091 |
| 6ayf | 7019 | 3.6 | FAT | **Input** | 17.90 | 1748 | 128 | 4 | 0 | 0.11% | 0.8066 | 0 | -22033 |
| | | | | **LF** | 1.11 | 1751 | 118 | 11 | 96 | 4.37% | 0.7782 | 0 | -23433 |
| | | | | **LF_RSR** | 2.83 | 1798 | 82 | 0 | 0 | 0.00% | 0.8105 | 0 | -27527 |
| 5a63 | 3061 | 3.4 | FAT | **Input** | 39.82 | 1108 | 77 | 26 | 8 | 13.74% | 0.8001 | 0 | -30726 |
| | | | | **LF** | 2.07 | 1095 | 82 | 34 | 83 | 5.44% | 0.7557 | 0 | -35101 |
| | | | | **LF_RSR** | 5.80 | 1123 | 87 | 1 | 0 | 0.67% | 0.8054 | 0 | -39137 |
| 5a1a | 2984 | 2.2 | FAT | **Input** | 11.62 | 3896 | 160 | 24 | 0 | 1.03% | 0.7636 | 56 | -240900 |
| | | | | **LF** | 1.53 | 3841 | 206 | 33 | 167 | 3.78% | 0.7364 | 56 | -232436 |
| | | | | **LF_RSR** | 5.51 | 3939 | 141 | 0 | 0 | 0.11% | 0.7694 | 56 | -251012 |
| 5kne | 8276 | 5.6 | CA | **Input** | 41.62 | 3221 | 656 | 83 | 0 | 0.00% | 0.8062 | 0 | 65030 |
| | | | | **LF** | 3.61 | 3128 | 636 | 196 | 269 | 5.93% | 0.7378 | 0 | -22207 |
| | | | | **LF_RSR** | 20.13 | 3437 | 515 | 8 | 0 | 0.46% | 0.8122 | 0 | -24327 |
| 5of4 | 3802 | 4.4 | FAT/PA | **Input** | 9.85 | 2079 | 265 | 15 | 20 | 1.90% | 0.6930 | 2 | -9493 |
| | | | | **LF** | 1.59 | 2104 | 172 | 83 | 120 | 5.07% | 0.6402 | 2 | -15031 |
| | | | | **LF_RSR** | 10.62 | 2115 | 244 | 0 | 0 | 0.40% | 0.7076 | 2 | -15839 |
| 5uj9 | 8559 | 3.5 | FAT/PA | **Input** | 0.65 | 1224 | 53 | 0 | 0 | 0.00% | 0.8129 | 0 | -42877 |
| | | | | **LF** | 1.05 | 1196 | 63 | 18 | 69 | 3.92% | 0.7937 | 0 | -39510 |
| | | | | **LF_RSR** | 3.51 | 1223 | 54 | 0 | 0 | 0.20% | 0.8274 | 0 | -44563 |
| 5u0p | 8479 | 4.4 | FAT/PA | **Input** | 17.73 | 2538 | 216 | 87 | 1 | 0.00% | 0.7496 | 42 | 3061 |
| | | | | **LF** | 3.14 | 2508 | 225 | 108 | 166 | 4.26% | 0.7543 | 42 | -28167 |
| | | | | **LF_RSR** | 11.35 | 2558 | 278 | 5 | 0 | 0.25% | 0.7703 | 42 | -29838 |

**Supplementary Table 1 (Continued)**

| pdb | map | res. (Å) | model type | | Clashscore | Ramachandran plot Favored | Allowed | Outliers | C-beta deviations | Rotamer outliers | phenix CC | Cis-peptides | Rosetta score |
|---|---|---|---|---|---|---|---|---|---|---|---|---|---|
| 5gar | 8016 | 6.4 | CA | **Input** | 0.00 | 5115 | 524 | 181 | 0 | 0.00% | 0.7404 | 6 | -38687 |
| | | | | **LF** | 4.30 | 5064 | 506 | 246 | 322 | 6.34% | 0.7670 | 6 | -56362 |
| | | | | **LF_RSR** | 10.37 | 5266 | 527 | 27 | 1 | 0.74% | 0.7621 | 6 | -63457 |
| 5uar | 8461 | 3.7 | FAT | **Input** | 0.26 | 1086 | 87 | 1 | 0 | 1.06% | 0.7837 | 0 | -16845 |
| | | | | **LF** | 0.94 | 1080 | 76 | 18 | 69 | 3.57% | 0.7521 | 0 | -14553 |
| | | | | **LF_RSR** | 4.84 | 1106 | 67 | 1 | 0 | 0.19% | 0.8074 | 0 | -17206 |
| 3j7h | 5995 | 3.2 | FAT | **Input** | 130.65 | 3972 | 100 | 8 | 0 | 12.24% | 0.8210 | 64 | 21515 |
| | | | | **LF** | 1.84 | 3774 | 235 | 71 | 204 | 5.98% | 0.8124 | 64 | -146830 |
| | | | | **LF_RSR** | 5.42 | 3864 | 208 | 8 | 0 | 0.11% | 0.8479 | 64 | -163189 |
| 5m54 | 4156 | 8.0 | FAT | **Input** | 7.01 | 1749 | 62 | 0 | 0 | 0.00% | 0.7978 | 2 | -26988 |
| | | | | **LF** | 1.63 | 1693 | 85 | 33 | 64 | 3.01% | 0.7763 | 2 | -24993 |
| | | | | **LF_RSR** | 8.31 | 1745 | 64 | 2 | 0 | 0.00% | 0.8306 | 2 | -27249 |
| 6bqr | 7132 | 3.2 | FAT/PA | **Input** | 25.51 | 3360 | 360 | 8 | 0 | 2.07% | 0.7606 | 0 | -76190 |
| | | | | **LF** | 1.82 | 3375 | 274 | 79 | 173 | 6.38% | 0.7529 | 0 | -105226 |
| | | | | **LF_RSR** | 5.03 | 3416 | 304 | 8 | 0 | 0.63% | 0.7946 | 0 | -122613 |
| 5vou | 8722 | 6.4 | FAT/PA | **Input** | 15.37 | 2132 | 134 | 0 | 0 | 0.00% | 0.8029 | 12 | 19334 |
| | | | | **LF** | 4.51 | 2103 | 128 | 35 | 110 | 4.29% | 0.7620 | 12 | -21193 |
| | | | | **LF_RSR** | 15.80 | 2079 | 186 | 1 | 0 | 0.52% | 0.8383 | 12 | -22414 |
| 5vkq | 8702 | 3.6 | FAT/PA | **Input** | 7.54 | 5428 | 532 | 20 | 0 | 0.72% | 0.7708 | 0 | -nan |
| | | | | **LF** | 3.33 | 5340 | 436 | 196 | 298 | 4.64% | 0.7610 | 0 | -60978 |
| | | | | **LF_RSR** | 6.90 | 5502 | 470 | 8 | 0 | 0.48% | 0.8133 | 0 | -76415 |
| 5oej | 3790 | 5.7 | PA | **Input** | 13.40 | 2264 | 240 | 115 | 0 | 0.00% | 0.5559 | 20 | 160946 |
| | | | | **LF** | 4.39 | 2318 | 174 | 127 | 135 | 6.07% | 0.4661 | 20 | -2014 |
| | | | | **LF_RSR** | 10.44 | 2308 | 296 | 15 | 0 | 0.32% | 0.6002 | 20 | -7555 |
| 5u1d | 8482 | 4.0 | FAT/PA | **Input** | 0.14 | 1090 | 66 | 2 | 0 | 0.84% | 0.7126 | 0 | 4463 |
| | | | | **LF** | 3.06 | 1057 | 74 | 27 | 58 | 3.73% | 0.6988 | 0 | -10391 |
| | | | | **LF_RSR** | 6.28 | 1087 | 70 | 1 | 0 | 0.10% | 0.7591 | 0 | -13015 |
| 5gw5 | 9541 | 4.6 | FAT | **Input** | 5.65 | 8025 | 382 | 7 | 0 | 0.01% | 0.6374 | 0 | -75861 |
| | | | | **LF** | 1.11 | 7842 | 452 | 120 | 320 | 2.43% | 0.6493 | 0 | -64126 |
| | | | | **LF_RSR** | 3.19 | 8059 | 339 | 16 | 0 | 0.03% | 0.6997 | 0 | -72841 |
| 3jac | 6343 | 4.8 | FAT/PA | **Input** | 59.58 | 2408 | 132 | 142 | 3 | 4.17% | 0.5443 | 10 | 78873 |
| | | | | **LF** | 4.55 | 2404 | 162 | 116 | 173 | 5.46% | 0.5659 | 10 | -29172 |
| | | | | **LF_RSR** | 11.01 | 2351 | 325 | 6 | 0 | 0.00% | 0.6105 | 9 | -30404 |
| 5ljo | 4061 | 4.9 | FAT | **Input** | 4.65 | 1401 | 151 | 62 | 1 | 0.15% | 0.7749 | 5 | -26521 |
| | | | | **LF** | 1.48 | 1412 | 154 | 48 | 62 | 3.01% | 0.8023 | 5 | -21902 |
| | | | | **LF_RSR** | 22.93 | 1432 | 180 | 2 | 0 | 0.15% | 0.8446 | 0 | -21066 |
| 5l93 | 4015 | 3.9 | FAT | **Input** | 3.37 | 601 | 65 | 0 | 0 | 0.00% | 0.7419 | 6 | -8106 |
| | | | | **LF** | 1.35 | 609 | 53 | 4 | 40 | 3.57% | 0.7657 | 6 | -7533 |
| | | | | **LF_RSR** | 4.71 | 633 | 33 | 0 | 0 | 0.00% | 0.8115 | 6 | -9163 |
| 5fxh | 3353 | 5.0 | PA | **Input** | 17.08 | 2730 | 277 | 14 | 0 | 0.00% | 0.7489 | 2 | 150477 |
| | | | | **LF** | 8.94 | 2585 | 297 | 131 | 215 | 6.70% | 0.7799 | 2 | -41193 |
| | | | | **LF_RSR** | 14.98 | 2739 | 273 | 9 | 3 | 0.79% | 0.8209 | 2 | -47853 |
| 3jd8 | 6640 | 4.4 | FAT/PA | **Input** | 75.57 | 897 | 134 | 94 | 5 | 11.82% | 0.6446 | 8 | 46603 |
| | | | | **LF** | 6.33 | 961 | 111 | 53 | 70 | 4.07% | 0.6560 | 8 | -6565 |
| | | | | **LF_RSR** | 9.86 | 986 | 135 | 4 | 0 | 0.31% | 0.7292 | 8 | -8114 |
| 3jc7 | 6536 | 4.8 | FAT | **Input** | 42.20 | 4479 | 424 | 66 | 7 | 0.45% | 0.6584 | 195 | 47295 |
| | | | | **LF** | 1.87 | 4307 | 472 | 196 | 319 | 5.81% | 0.6867 | 196 | -30588 |
| | | | | **LF_RSR** | 11.57 | 4395 | 557 | 23 | 2 | 0.31% | 0.7509 | 192 | -34889 |
| 3jch | 6553 | 7.1 | CA | **Input** | 2.28 | 1567 | 80 | 0 | 0 | 0.00% | 0.7030 | 0 | -20566 |
| | | | | **LF** | 5.10 | 1517 | 79 | 51 | 87 | 5.49% | 0.7791 | 0 | -20090 |
| | | | | **LF_RSR** | 14.00 | 1526 | 121 | 0 | 0 | 0.06% | 0.8489 | 0 | -23141 |
| 6eny | 3906 | 5.8 | PA | **Input** | 12.90 | 1368 | 136 | 33 | 0 | 0.00% | 0.5648 | 12 | 61616 |
| | | | | **LF** | 7.94 | 1312 | 163 | 64 | 71 | 6.19% | 0.7795 | 12 | -12663 |
| | | | | **LF_RSR** | 17.71 | 1383 | 154 | 2 | 0 | 0.60% | 0.7362 | 12 | -15692 |
| 5nd7 | 3623 | 7.9 | FAT | **Input** | 19.97 | 1024 | 84 | 10 | 6 | 14.15% | 0.5818 | 0 | -10039 |
| | | | | **LF** | 1.57 | 998 | 89 | 31 | 62 | 5.91% | 0.6993 | 0 | -12488 |
| | | | | **LF_RSR** | 8.10 | 1034 | 83 | 1 | 0 | 0.00% | 0.7673 | 0 | -13362 |
